# Accuracy of somatic variant detection workflows for whole genome sequencing experiments

**DOI:** 10.1101/2021.06.10.446467

**Authors:** Roman Jaksik, Jacek Rosiak, Paweł Zawadzki, Paweł Sztromwasser

## Abstract

Whole genome sequencing (WGS) becomes increasingly important for advancing personalized cancer care, driving not only basic science studies but also entering into clinical applications. Translating raw WGS data into the right clinical decision requires high accuracy of somatic variant detection, therefore novel data analysis methods have to be carefully evaluated.

In this work we tested the performance of well-established somatic variant detection workflows: GATK, CPG-WGS, DRAGEN and Strelka2. By utilizing both real data, with well-defined mutations, and synthetic mutations spiked-in into real data, we were able to assess sensitivity and precision of each workflow, for various coverage and tumor purity levels.

Individual tools excelled in different evaluation approaches, however the results demonstrated that DRAGEN has the highest overall performance when sensitivity is preferred over precision, and the opposite is true for CGP-WGS. The differences in results obtained using synthetic and real datasets, indicate that benchmarks based only on a single reference set may provide an incomplete picture.

## Introduction

Next-generation sequencing (NGS) technologies have become invaluable for the development of cancer therapies, driving not only the basic science studies [1, 2] but also translating to clinical applications, influencing the selection of best treatment strategy [3]. DNA sequencing, based on short-read approaches, have been widely used for the detection of somatic variants, which allow to determine tumor driving genomic alterations and changes that help to predict the effectiveness of anticancer therapies [4, 5]. It is therefore important that DNA sequencing provides accurate information which will translate to the right clinical decision. Financial constraints favor the use of gene-panel or whole exome sequencing (WES) in studies focusing only on short variants, located in the coding regions [6], however, whole genome sequencing (WGS) is superior by providing unbiased and uniform read coverage, insights into variants located in intronic and intergenic regions, greater sensitivity in detecting structural variants and copy number changes [7].

While the decreasing costs of whole genome-sequencing (WGS) experiments gradually make it applicable to characterize the mutational landscape of tumors on a routine basis [8], the precision of variant discovery methods are still insufficient to rely on the provided results without a proper validation [9], capabilities of which are significantly limited not only by the costs of validating a single variant but by the dependence on similar error prone DNA handling protocols [10]. The accuracy of somatic variant detection methods depends on their ability to deal with current technological limitations associated with NGS approaches, including uneven coverage [7], sequencing error rate [11], strand bias [12], deamination artifacts [13], PCR amplification errors [14] and incorrect local alignments of reads [15]. Tumor heterogeneity and clonality leading to multiallelic variants or low variant allele frequency levels (VAF) [16], DNA copy number aberrations, as well as contaminants in both tumor and reference sample [17, 18], make this process even more challenging.

The problem of somatic variant discovery was addressed many times by the scientific community resulting in a high number of dedicated tools, which were designed using various statistical models or machine learning approaches [19]. The choice of an individual tool, which would deliver the most accurate detection results is extremely difficult, due lack of high quality gold standards. In contrast to a number of reference germline genomes [20] no tumor genome has been fully characterized for the purpose of benchmarking variant discovery methods. The most typical approach to developing a validated truth-set of reference variants is to confirm the NGS-derived variant calls using target DNA capture, followed by high coverage NGS approaches (e.g. Ampli-seq), RT-qPCR, Sanger sequencing [21-23], or manual curation [24], using read visualization tools [25, 26]. However, while this is a viable method of validating individual variants it is not the best choice for the development of a gold standard set, for the purpose of comparing variant callers. The main limitation results from the fact that the validation will include only those variants which were successfully detected in the NGS experiment, using one or few variant callers in the process, and at the same time biasing the benchmarking results towards them. Another limitation is associated with the fact that the validation methods are not applicable for all variants e.g. Sanger sequencing is not suited for the detection of low VAF variants [27]. The inability to validate all detected positions due to technical reasons of financial constraints can lead to underestimated true-positive (TP) rate and overestimated false-positive (FP) rate potentially biasing the results towards methods that favor precision above sensitivity.

Another approach is to derive a gold standard based on concordance between multiple callers [28-30], assuming that variants identified only by a single caller are likely false positives. The main limitation of this approach is that methods based on similar assumptions are likely to have a leading vote on what is an actual variant. Additionally the tools used to create the gold standard will have an advantage above the rest, which potentially biases the results of a benchmarking experiment.

Third possibility is to utilize synthetic data generated by either mixing two samples, with known germline variants in various proportions [31-33], by introducing variants into existing reads [34, 35], or by generating synthetic reads [36]. The main advantage of this approach is that the dataset creation results in the exact coordinates of all variants. The main weakness is that such synthetic dataset might not well represent a typical cancer genome due to sequencing error rate of artificial characteristics, unrealistic distribution of VAF, lack or unrealistic structural variants and copy number events, or other potential factors which we are unaware of, that affect the detection accuracy.

Different strategies to the creation of gold standards were adopted in existing benchmarking studies, leading to significantly different outcomes, which in many cases lack clear recommendations. The majority of benchmarking studies focus on exome or amplicon sequencing [22-24, 31, 32, 37, 38], which provides high but uneven coverage, biased by the target enrichment strategies utilized. Only few studies focus on whole genome sequencing, using a gold standard originating from PCR/Sanger validation [21], gold standard based on method concordance [39, 40], or synthetic data [40]. The number of compared tools is usually limited to only the most popular approaches, with the total number per study varying between two [37] and nine [24], however most studies focus on four callers only, testing their performance using the default parameters.

In this work we aimed to assess the accuracy of methods used to identify somatic single nucleotide variants (SNVs) and short insertions/deletions (indels), specifically in WGS experiments. For this purpose we used two distinct truth sets based on concordance between multiple detection methods as described in [30], and based on synthetic variants inserted into reads obtained by sequencing a germline genome [20]. We focused not on accuracy of individual somatic callers but rather well-established workflows: GATK (based on Mutect2) [41] developed at the Broad Institute; CGP-WGS (based on CaVEMan [42] and Pindel [43]) developed by Cancer IT at Sanger; Dynamic Read Analysis for GENomics (DRAGEN) [44], a platform developed by Illumina, which was shown to have high accuracy of germline variant detection [45]; and standalone Strelka2 [46] variant caller, also developed by Illumina. The main goals of this work were therefore 1) to evaluate the performance of each workflow; 2) compare the outcomes between the two selected gold standards; 3) determine the characteristics of method-specific variants.

## Results and discussion

### Selection of datasets

Due to the limitations of methods used to evaluate accuracy of variant detection we decided to include three distinct approaches. The first one is based on data provided by the Somatic Working Group of SEQC-II Consortium, described in the Fang et al. article [30], which is based on the truth set generated using multiple high coverage sequencing replicates and various combinations of data analysis algorithms, including six variant callers. We used the high confidence set which comprises 39632 SNVs and 1939 indels. We refer to this dataset as Fang2019 in this manuscript. The second dataset was created by inserting randomly generated variants into reads of the HG001 genome, thoroughly studied by the Genome in a Bottle Consortium [20]. In total we inserted 16995 SNVs and 1700 indels (see materials and methods for details). We refer to this dataset as Synthetic in this manuscript. The third dataset was based on reads obtained purely from normal tissues which we used in normal-to-normal sample analyses, to evaluate the false positive rate of each workflow.

The datasets were used to test the association between sequencing coverage and cancer sample purity on the sensitivity and precision of both SNVs and indel detection, using 4 data analysis workflows, summarized on Fig.1. Additionally, we included in the comparison a tumor-only analysis offered by the DRAGEN pipeline (not shown in Fig.1). In total, we analyzed 63 SNV and indel callsets.

**Fig.1:**
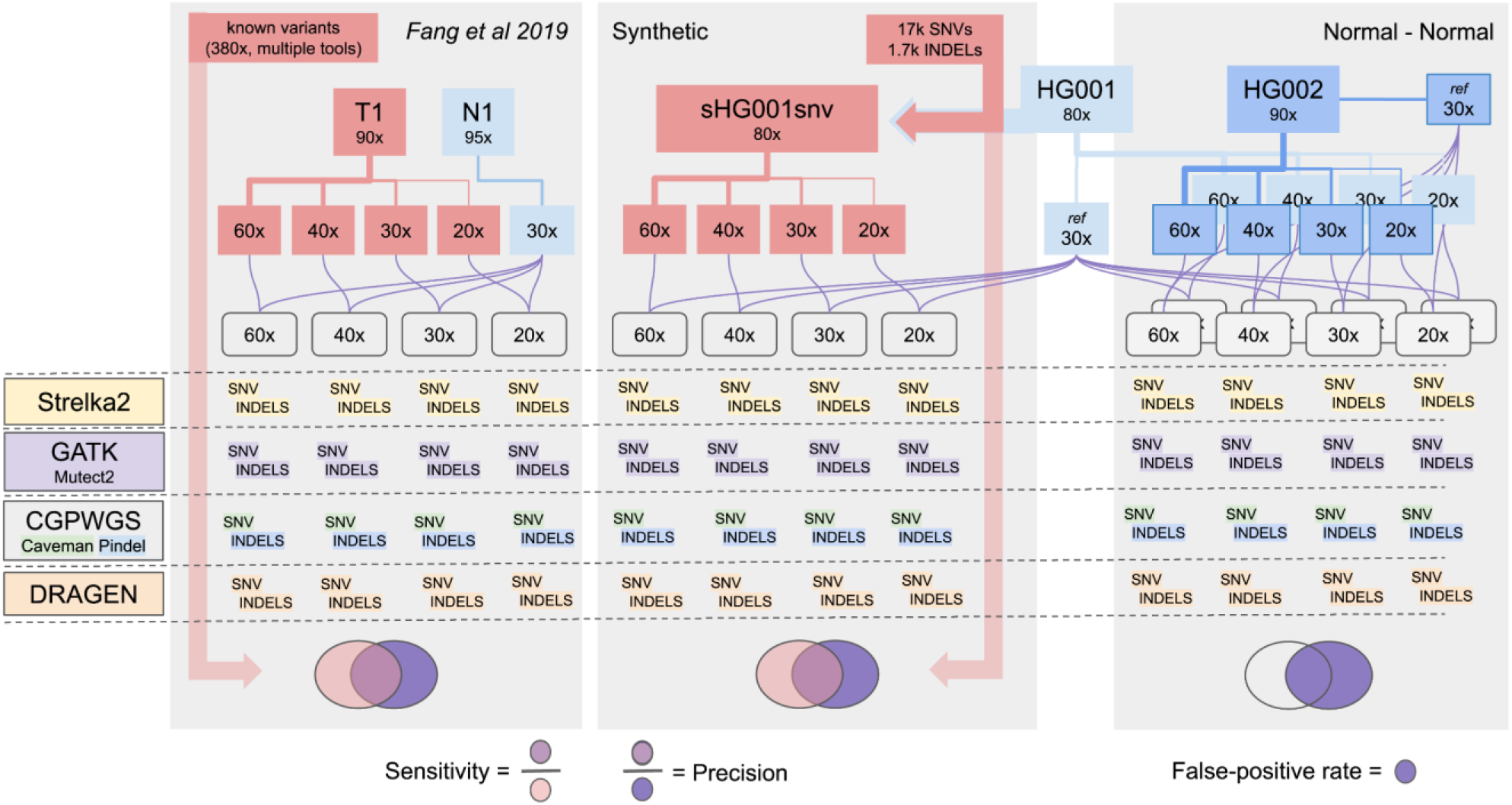
Overview of the analysis methods and dataset utilized in this study

### Accuracy of somatic variant detection

The performance of all utilized somatic variant detection workflows was evaluated using sensitivity and precision metrics obtained by comparing the calls to corresponding truth sets. The comparisons were carried out for four coverage levels of the tumor sample, ranging from 20x to 60x, and four tumor sample purity levels.

Fig. 2A shows the association between sensitivity and precision for various coverage levels of the tumor sample, separately for SNVs and indels. The differences in performance levels of individual workflows between both datasets were substantial and variability between workflows for individual datasets considerably higher in the Fang2019 compared to the Synthetic set. It is worth noting that the overall precision of SNV detection in the Synthetic dataset was higher than 0.8, while analyses on Fang2019 showed a generally higher sensitivity. In all tests precision was only marginally affected by the reduced coverage levels, while sensitivity was more significantly affected in the Synthetic dataset. This likely results from a higher number of variants with low VAFs (see further on).

**Fig. 2:**
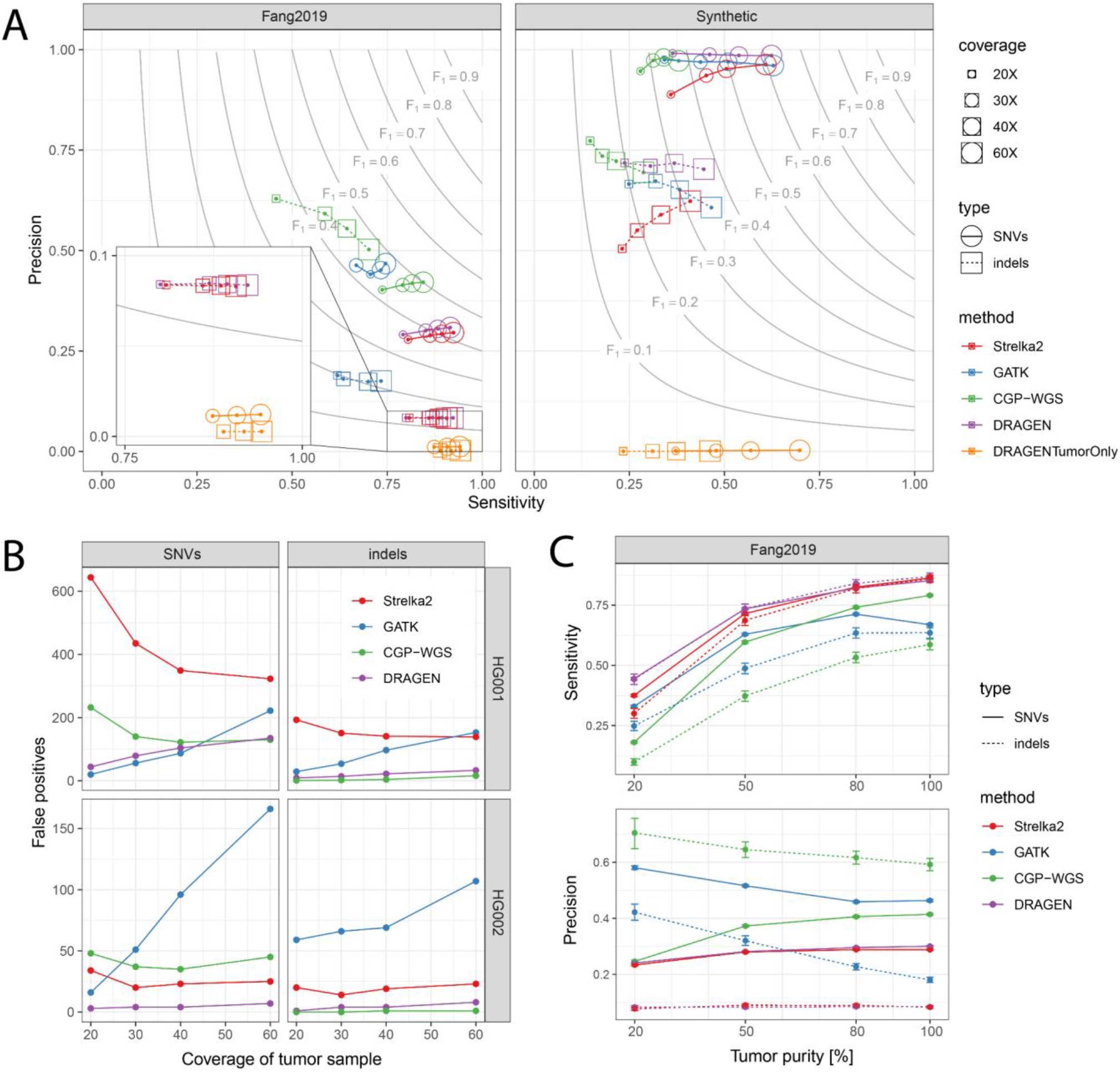
Performance of variant detection workflows on real and synthetic data: A) Precision and sensitivity of SNV and indel detection obtained for various coverage levels; Gray lines mark specific F1 statistic values B) Total number of false positives obtained by comparing two replicates of GIAB samples (HG001 and HG002); C) Sensitivity and precision of variant detection obtained for different cancer purity levels.

On the Fang2019 dataset GATK showed the lowest sensitivity but highest precision of SNV detection, despite variable coverage levels. However, on the Synthetic dataset GATK showed the highest SNV sensitivity for 60X coverage but performed worse for lower coverage levels, although the differences were not substantial. Surprisingly, the precision of GATK at 30x coverage in the Fang2019 set was worse compared to 20x coverage. Similar trend was not observable for other workflows.

Comparable performance metrics, obtained by the DRAGEN and Strelka2 pipelines for the Fang2019 dataset, indicates that the two tools developed by Illumina share some similarity. This concordance in accuracy disappears in the results obtained for the Synthetic set where DRAGEN clearly outperforms Strelka2, in both indel and SNV detection for lower coverages (20-40), indicating that the pipeline is better tuned for detection of low frequency variants. Generally, DRAGEN achieved the highest precision and sensitivity for SNVs and indels in the Synthetic dataset (highest F1 score), showing only marginally worse sensitivity at 60x coverage and smaller F1 for indels at 20x and 30x coverage levels, compared to GATK.

DRAGENTumorOnly was an outlier in all comparisons. It showed the highest sensitivity among all methods, but due to absence of the normal/reference sample, the pipeline was not able to discriminate a large portion of germline variants, resulting in very low precision in all tests (on average 0.006 for SNVs and 0.002 for indels). To better capture somatic variation, results of this pipeline would need additional filtering based on e.g. population frequency of the variants.

CGP-WGS, which utilizes CaVEMan for SNV detection and Pindel for indels, showed high precision in both datasets but at the cost of low sensitivity. This is especially evident for indel detection where the differences in the numbers of identified variants between all methods were substantial. CGP-WGS provided 2707 indels at 60x coverage in Fang2019 set, which is three times less compared to GATK, and eight times less compared to Strelka2 and DRAGEN. In the synthetic set CGP-WGS also provided the smallest number of indels, however the differences were not as large, since at the 60x coverage the numbers of indels identified by individual methods ranged between 701 and 1301. Detailed results, used to create Fig.2A, are available in the supplementary table S1.

Fig.2B shows a different workflow evaluation approach (Normal-normal on Fig.1), which was based on false positive (FP) rates obtained by comparing two subsets of reads of the same sample. The comparison was performed on two samples, HG001 (GM12878) and HG002 (GM24385), and while the differences between both samples are substantial some trends are evident. For the GATK, the false positive rate increases significantly with the increasing tumor coverage level. This trend for the GATK was also observed on Fig.2A where the precision decreased with increasing tumor coverage. The reverse trend was observed in the case of Strelka2 - the increasing coverage of the tumor sample allowed to reduce the FP-rate. Interestingly, while GATK showed low FP-rates for the HG001 genome, and highest in case of HG002, Strelka2 performed well in the HG002 and worst in HG001. In general the lowest FP rate, largely unaffected by the coverage levels, was observed for the DRAGEN pipeline. DRAGEN showed the best performance for SNVs and was only marginally worse for indels compared to CGP-WGS. For detailed results see Supplementary Table S2.

Significant differences between results obtained using the Synthetic and Fang2019 datasets likely stem from different assumptions made during the creation of the gold standard, but also from the specificity of the samples themselves. Fang2019 truth set was created based on concordance between multiple methods, including six mutation callers (MuTect2 [33], SomaticSniper [47], VarDict [48], MuSE [49], Strelka2 [46], TNscope [50]). It is therefore potentially biased towards MuTect2, Strelka2, and possibly DRAGEN used in our comparison. The Synthetic dataset compensates this potential bias as it was created independently of variant detection algorithms. Additionally, while Fang2019 dataset included only high confidence variants with relatively high average VAF, which are expected to be more easily detected, it also included a very high number of copy number alterations and loss of heterozygosity regions (data not shown) making the detection more difficult. The Synthetic dataset, although based on a more realistic distribution of VAFs observable in cancer genomes, did not contain any copy number changes, nor structural variants frequently found in tumors. It is also worth noting that, while the read sampling for low coverage levels was conducted in the same way for the Fang2019 and the Synthetic sets, the true coverage obtained after read processing differed substantially (supplementary figure S1A). While the fraction duplicate reads was roughly 2x higher in the Synthetic set leading to higher data loss (Fig. S1C), the insert size distribution in this set had a higher mean resulting in lower data loss caused by read overlaps in paired-end sequencing (Fig. S1B). Also, the coverage uniformity in the Synthetic set is much higher compared to the Fang2019 dataset, leading to the higher median coverage across all genomic positions.

### Tumor sample purity

Contamination of the tumor sample with normal cells is a common problem which affects variant detection sensitivity resulting from a lower number of reads that support the existence of a particular variant [32]. To test the influence of tumor sample purity on the detection accuracy we created synthetic dilutions of the tumor sample by mixing-in normal sample reads to the tumor in various proportions, resulting in 20, 50 and 80% purity levels.

Fig.2C shows the sensitivity and precision obtained for various tumor purity levels of Fang2019 at 30x coverage (for detailed results see Supplementary Table S3). Not surprisingly tumor purity has the highest impact on the sensitivity of variant detection. The results indicate that sensitivity of all tools is similarly affected by reduced tumor purity, and all tools perform relatively well for 50% and higher purity. The impact on precision was more variable, although we observed that precision of indel detection was more significantly affected by the tumor purity, than SNV detection. Changes in the purity levels generally did not have a significant impact on the relative performance of each tool. An exception is CGP-WGS which was more sensitive for high purity samples (80% and 100%), and showed lower sensitivity in more contaminated samples (20 and 50%), when compared to GATK. An interesting fact is that precision of GATK drops as the purity increases, a trend which can be also observed only for the indels identified using CGP-WGS (Pindel). Notably, sensitivity of GATK drops at 100% purity compared to 80%, which results from a 15% higher number of FN and slightly lower TP rate. For the 80% purity GATK identified 4326 more variants compared to the 100% purity sample.

### VAF-based filtering

Precision of variant detection is expected to be affected by the variant allele frequency (VAF). The higher the VAF of a variant, the higher is the detection probability, since more potential evidence is available in a form of reads that support the modified allele. Also it is expected that the higher the VAF of the identified variant the lower the probability that a particular site is a false positive resulting from a sequencing error. Therefore, in theory, it should be easier to detect variants with high VAF. By focusing the performance evaluation on variants with high VAF, the FP- and FN-rates should decrease (compared to all-variants experiments) resulting in higher precision and sensitivity values.

Fig. 3A shows the distribution of VAF of all gold standard variants from both datasets. Fang2019 variants are pre-filtered by their authors, which is likely why low VAF variants are underrepresented in this set. It also includes a high number of variants with VAF∼1, resulting from a large fraction of loss of heterozygosity regions in this cell-line sample. Distribution of VAFs in the Synthetic set reflects a distribution typically observed in tumor samples [51, 52], resulting from their subclonal architecture (in the absence of subclonal selection). Due to the high number of variants supported by a small number of reads, the detection process in this set should be more challenging, or even impossible in sites with insufficient read depth (samples with coverage reduced below 60x).

**Fig. 3:**
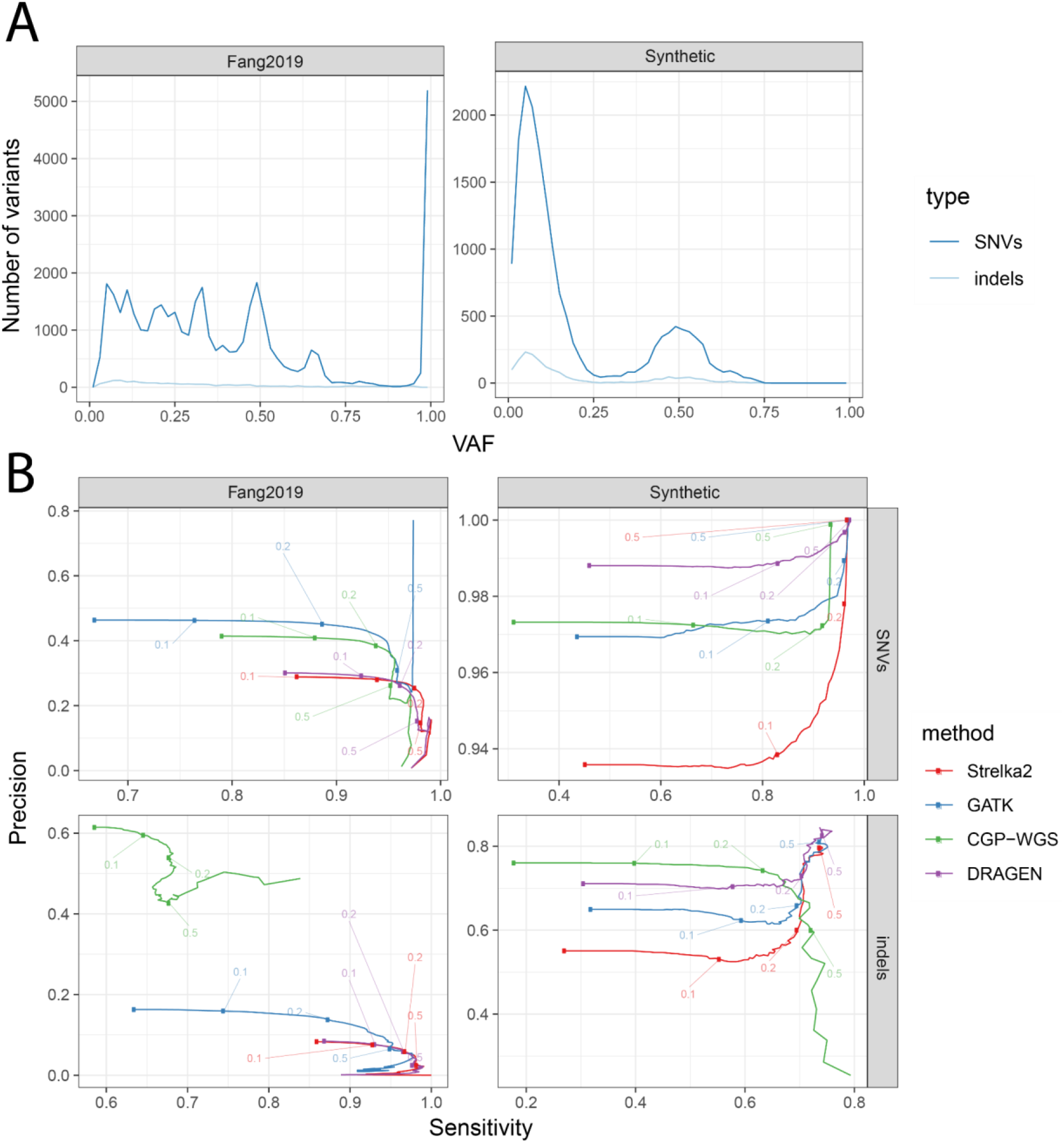
Effects of VAF-based variant filtration on the accuracy of variant detection: A) distributions of VAF estimates obtained for SNVs and indels in both datasets used; B) Precision–Sensitivity curves obtained for various VAF cutoff levels

Fig. 3B shows the association between precision and sensitivity for various VAF cutoff levels (only variants with VAF higher than the threshold are used to calculate both metrics). The filtering is carried out based on expected VAFs (using gold standard) for TP and FN variants and based on those provided by the callers for FP variants. VAF based filtering in the Fang2019 dataset leads to a reduced sensitivity and precision, which is counter intuitive. This results from a high number of FP variants with high VAF, which might be associated with the way the variants were selected for the high confidence set. The same does not apply to the Synthetic set where in most cases sensitivity and precision obtained for variants with VAF>0.1 was substantially higher. Pindel is an exception from this rule, which may be associated with strict variant filtering criteria in the CGP-WGS pipeline, resulting in a significantly lower number of identified indels (Fig. 2).

### Characteristics of method-specific variants

The lack of concordance between variant detection workflows results not only from various precision and recall levels but also from different filtering strategies that are utilized as a part of the post processing steps. For this reason it is important to assess what are the characteristics of variants identified by a single method and by the majority of methods.

Fig. 4 shows the number of variants identified by a specific group of workflows in both datasets and separately for SNVs and indels. In all cases VAF distribution of variants identified by all methods has a median in the vicinity of 0.5 representing a group which is likely the easiest to identify, due to the potentially relatively large number of reads that support them. Also, the variants which were not detected by any method have the lowest median VAF among all compared groups. Interestingly the variants which were not detected only by the CPG-WGS workflow in the Fang2019 set have a VAF distribution similar to that observable for variants identified by all methods, with relatively high median. This could be due to the overly strict filtering employed in the CGP-WGS pipeline tuned for high precision.

**Fig. 4:**
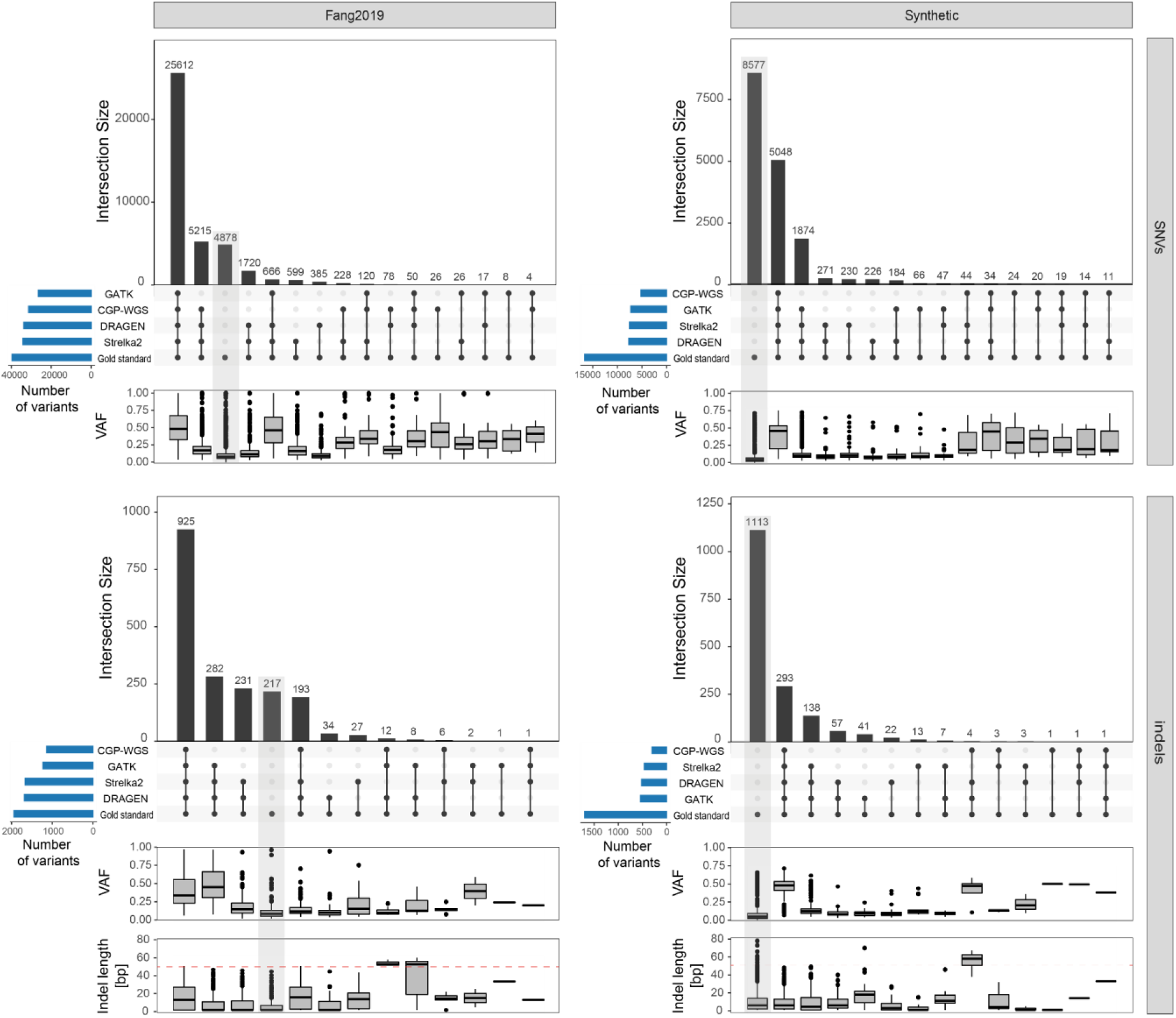
Total number, VAF distribution and indel length statistics, of true variants identified using a specific subset of workflows for tumor samples with 30X coverage. Light gray bars highlight variants which were not detected by any of the methods utilized (observed in the gold standard only). Boxplots below each of the bar plots show the distribution of VAFs and indel lengths for each of the variant groups. Red dashed line on the indel length plot marks the 50bp cutoff.

In both sets there are very few variants which were detected by only a single method, 2.6% and 3.2% for SNVs and indels respectively in the Fang2019 dataset, and 1.9% and 4.5% in the Synthetic set. In both datasets Strelka2 contributed the most to the percentage for SNVs by providing the highest number of variants which were not detected by any other method, with DRAGEN taking the second position. For indels the results differ between both sets, for Fang2019 the highest number of method specific variants were provided again by Strelka2 and DRAGEN, however in the Synthetic set, GATK took the lead.

In both datasets, indels not detected by Strelka2 but found by all other methods, are all longer than 50nt. This is the result of the maximum indel length which in Strelka2 is set to strict 50nt. The constraint is not present in the DRAGEN pipeline which, although showing similar performance, was not affected by this limitation.

## Summary

In this work we tested the performance of well-established somatic variant detection workflows GATK [41], CPG-WGS [42, 43], DRAGEN [44], and standalone Strelka2 [46], using several evaluation approaches. Estimates of sensitivity and precision, for various coverage and tumor purity levels, were obtained using two reference datasets: the Fang2019 gold standard derived using concordance between multiple detection tools [30], and a synthetic dataset generated by spiking-in randomly generated variants to real sequencing data from a well characterized HG001 genome [20]. We also investigated the impact of tumor sequencing depth on the variant calling false positive rate by comparing two pairs of well-known germline reference samples [20]. While the results obtained using different approaches showed significant differences, favoring various tools at specific categories, in our opinion DRAGEN showed the highest overall performance when favoring sensitivity above precision, and CGP-WGS when precision is of higher importance. In the Synthetic dataset DRAGEN showed the highest precision among all tools, both for SNVs and indels and only marginally worse sensitivity compared to GATK. In the Fang2019 dataset it showed significantly lower precision, but the highest sensitivity for all tested coverage and tumor purity levels. DRAGEN also showed the overall lowest false positive rate in the comparison of two control samples when considering both SNVs and indels, although for indels alone it was slightly worse than CGP-WGS.

Inclusion of two reference datasets in our experiments allowed us to observe the impact of the truth set on the general conclusions regarding accuracy of the benchmarked tools, as well as their relative performance. While all tools showed similarly high precision, and comparable recall on the Synthetic dataset, Dragen and Strelka2 were significantly more sensitive, and less precise than GATK, when evaluated on the Fang2019 dataset. The overall sensitivity in Fang2019 was relatively high for all workflows, indicating that the top tools detect almost 90% of somatic variants. On the other hand, when compared on the Synthetic set, the sensitivity of variant detection was generally lower and precision was very high, suggesting that false-positive calls for SNV detection tools are a marginal problem. While both datasets have their own weaknesses we believe that by including them both in our study we were able to show the performance of all workflows in vastly different situations. Our results show that benchmarks based only on one dataset might be biased and methods fine-tuned on one dataset might perform badly on others.

The main weakness of our study, similarly to other benchmarking studies mentioned in the introduction, is that it focuses only on a single set of parameters of each tool, while fine tuning each method for optimal performance could potentially lead to better results. Most tools are highly parameterized allowing to affect the detection sensitivity and specificity levels, however the complexity of underlying statistical models and ad-hoc filters make them very difficult to safely manipulate by the scientists not involved in their development process. This makes the comparison process very difficult, limiting it to the default parameter set only, which in many cases are modified between individual releases resulting in significantly different accuracy levels. This was the main motivation behind focusing the study on established workflows, which as we assume, are already fine-tuned and represent a state in which the tool would be typically used by the end-user.

## Materials and methods

### Datasets used

#### HG001 and HG002

Raw WGS data for HG001 and HG002 genomes were downloaded from the Sequence Read Archive (SRA). For HG001, we combined data from multiple runs deposited by the Hartwig Medical Foundation in study ERP115966: ERR3607783, ERR3610392, ERR3610755, ERR3610808, ERR3582426, ERR3582714, ERR3584441, ERR3585465 (785.5M read-pairs). For HG002 we merged reads from 4 runs (SRR8861483, SRR8861484, SRR8861485, and SRR8861486; a total of 894.8M read-pairs) deposited by the Genome in a BottleConsortium [53] under experiment SRX5648942. In all cases the reads were paired-end, 2×150bp sequenced on Illumina Novaseq 6000 platform.

#### Fang2019

Fang *et al*. characterized a human triple-negative breast cancer cell line and a matched normal cell line as a reference set for somatic variants [30]. Illumina sequencing data deposited in the Sequence Read Archive under project SRP162370 available for each of the two samples exceeds 300x coverage. We selected two runs for the tumor T1 sample (SRR7890904, SRR7890905), hereafter and in Fig.1 referred to as T1, and two for the normal sample (SRR7890942, SRR7890943), referred to as N1. Combined runs had 1,245M and 1,387M trimmed 2×150bp read-pairs for the tumor and normal samples, respectively. Release 1.1 of the reference variants in the tumor sample, as well as high confidence regions used to limit the genomic regions considered in our analyzes, were downloaded from: https://ftp-trace.ncbi.nlm.nih.gov/seqc/ftp/Somatic_Mutation_WG/release/v1.1

### Generation of synthetic data

Paired-end FASTQ files for the HG001 sample were mapped to the GRCh37 reference genome as detailed in the following section. Next, BAMSurgeon v1.2 [35] was used to generate and spike-in small variants (point mutations and indels) into the HG001 BAM file. Generation of random variants was performed with the help of *randomsites*.*py* script from the BAMSurgeon framework. We simulated 3 subclones with varying parameters of variant allele fraction distributions: C1 (*--minvaf 0, --maxvaf 0*.*5, --vafbeta1 2, --vafbeta2 10*), C2 (*--maxvaf 0*.*75, --minvaf 0*.*25*), and C3 (*-maxvaf 0*.*6, --minvaf 0*.*4*). A total of 16,995 SNVs (C1:13000; C2:2000; C3:2000) and 1,700 indels (C1: 1300; C2: 200; C3: 200) were inserted into the reads within the high confidence regions as defined by the BED file in the 3.3.2 release of Genome-In-A-Bottle HG001 reference dataset (https://ftp-trace.ncbi.nlm.nih.gov/ReferenceSamples/giab/release/NA12878_HG001/NISTv3.3.2/GRCh37/HG001_GRCh37_GIAB_highconf_CG-IllFB-IllGATKHC-Ion-10X-SOLID_CHROM1-X_v.3.3.2_highconf_nosomaticdel.bed). In the end, reads from the resulting BAM file with mutation spike-ins, hereafter referred to as sHG001, were extracted using Picard v2.21.4 [54], and subjected to downsampling as shown in Fig.1

### Data pre-processing

Quality of the downloaded raw data was confirmed using FastQC v0.11.7 [55]. The full-size FASTQ files for tumor (sHG001 and T1) and normal samples (HG001, HG002, N1) were downsampled to approx. 20x, 30x, 40x and 60x coverage datasets using seqtk v1.3 tool (https://github.com/lh3/seqtk). Numbers of pseudo-randomly drawn read-pairs were 246.7, 370, 493.3, and 740M, for 20, 30, 40, and 60x respectively. The randomization seed was fixed for subsampling of each genome, so that lower coverage dataset was a subset of higher coverage dataset (e.g. all the read-pairs drawn for the 30x dataset were present in the 40x dataset).

The T1 sample derived from a cell line was 100% pure tumor tissue. To simulate more natural conditions we mixed reads from the T1 and N1 samples in 4:1, 1:1, and 1:4 proportions to obtain tumor purity levels of 80, 50, and 20%. Read sampling was performed as above to extract a total of 370M reads-pairs (∼30x) in each tumor dilution.

### Read mapping

All reads were mapped to the GRCh37 reference genome. Variant calls in the DRAGEN pipeline were obtained on alignments produced by the DRAGEN Alignment pipeline v.3.7.5. Alignments in all other pipelines were generated using the Sanger’s CGPMAP pipeline v3.0.0 (https://github.com/cancerit/dockstore-cgpmap), including BWA MEM [56] mapping, and alignment post processing with SAMtools [57] and biobambam2 [58]. Picard v2.18.26 [54] was used to produce mapping metrics.

### Variant detection

SNVs and indels were called on matched tumor and normal, as well as normal-normal samples using DRAGEN Somatic Analysis Pipeline v.3.7.5, CGP-WGS pipeline v2.0.1 (https://github.com/cancerit/dockstore-cgpwgs) developed by Cancer IT at the Wellcome Trust Sanger Institute, GATK somatic analysis pipeline v4.1.9.0 [41], and Strelka2 v.2.9.2 [46]. In addition, DRAGEN calls in tumor-only analysis (without normal sample) were generated. In all cases default parameters were used, and (unless noted otherwise) only variants with the PASS flag were used for the analysis. GATK pipeline was based on MuTect2, and the analysis was carried out according to the GATK recommendations: reads aligned as described above were processed using MarkDuplicates algorithm from the Picard tool set [54] and BaseRecalibrator which is a part of the Genome Analysis Toolkit (GATK v4.1.9.0) [41]. Somatic mutations were identified using MuTect2 (v4.1.9.0) [33] and filtered using GATK’s FilterMutectCalls, as well as sample contamination estimates obtained using CalculateContamination tool and read orientation bias statistics obtained with LearnReadOrientationModel tool.

### Accuracy of variant detection

Variant calls obtained with each of the tools were compared to the gold standard using the som.py tool from the hap.py package developed by Illumina [59]. The metrics used in the basic comparison, illustrated on Fig.1, were obtained by eliminating variants outside of the defined regions and without the PASS filtering flag. From the Synthetic set we used only variants identified in the autosomal and sex chromosomes. For the Fang2019 set only variants from the high confidence regions, as defined in the High-Confidence_Regions.bed file, were included in the study. Since the gold standard for Fang2019 set was based on hg38, prior to the comparison we converted all variant calls to GRCh37 using the liftOver tool provided by the UCSC [60] with the b37ToHg38.over.chain file. On average 1.85% of positions could not be converted due to location inside contig that existed only in the hg38 reference. While the percentage of unconverted positions differed between individual tools almost all were located outside of the high confidence regions defined in the Fang2019 set, and therefore would not be used as a part of the comparison, eliminating the conversion bias.

Precision-recall curves were obtained by running som.py with the --keep-scratch option which allowed us to extract individual variant positions that were later processed in R. The VAF cutoff levels used to create the curves were selected by dividing the VAF distribution into 0-90th percentiles. This allowed us to calculate the precision and sensitivity after iteratively excluding 1% of lowest VAF variants, making sure that the last point contained at least 10% variants of the entire set (reducing information noise). The filtering was carried out based on expected VAFs (defined in the gold standards) for TP and FN variants, and based on those provided by the callers for FP variants.

Fig. 4 was created using the UpSetR package [61] which shows a stratification of variants from the reference sets, along with information on which group of variants was identified by specific subset of workflows.

## Supporting information

Supplemental plot 1

## Acknowledgements

We would like to thank Illumina for providing computational credits for benchmarking the Illumina Dragen platform. Other calculations were performed at the Poznan Supercomputing and Networking Center (PSNC), and using infrastructure of the Ziemowit computer cluster (www.ziemowit.hpc.polsl.pl) in the Laboratory of Bioinformatics and Computational Biology, The Biotechnology, Bioengineering and Bioinformatics Centre Silesian BIO-FARMA, created in the POIG.02.01.00-00-166/08 and expanded in the POIG.02.03.01-00-040/13 projects. This work was supported by the Polish National Science Centre grant No. 2016/23/D/ST7/03665 for RJ and by the Foundation for Polish Science grant No. First TEAM/2016-1/9 for PZ.

