## Supplemental plot 1 for "Accuracy of somatic variant detection workflows for whole genome sequencing experiments"

### Supplement Plots

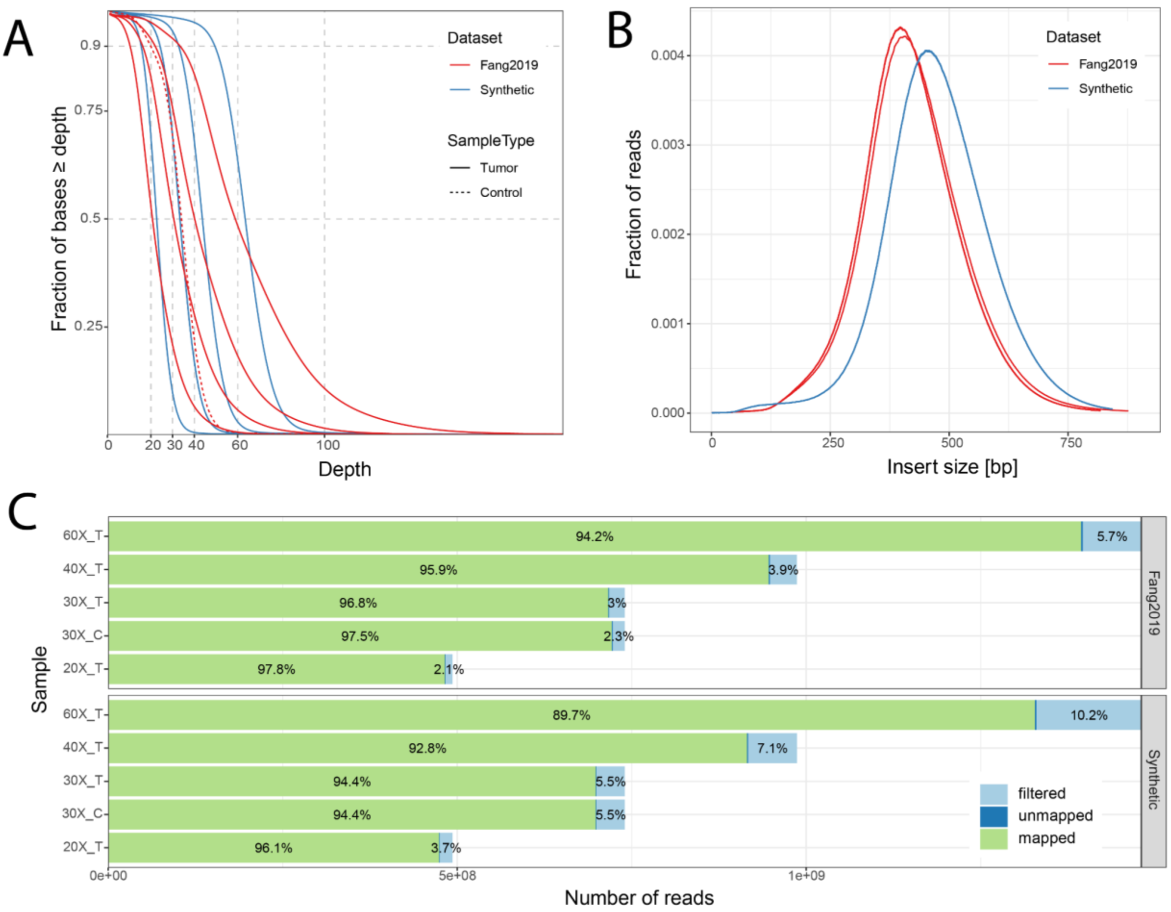


**Fig. S1**: Quality control statistics obtained for both datasets: **A)** fraction of genomic positions with coverage above specific threshold; **B)** Insert size distribution of individual samples from both utilized datasets; **C)** Read mapping and filtering statistics; the filtered group of reads includes low quality reads and duplicates. T and C suffixes in the sample names mark the tumor and control samples respectively.
